# Cytoplasmic nanojunctions between lysosomes and sarcoplasmic reticulum are required for specific calcium signaling

**DOI:** 10.1101/002196

**Authors:** Nicola Fameli, Oluseye A. Ogunbayo, Cornelis van Breemen, A. Mark Evans

## Abstract

Herein we demonstrate how nanojunctions between lysosomes and sarcoplasmic reticulum (L-SR junctions) serve to couple lysosomal activation to regenerative, ryanodine receptor-mediated cellular Ca^2+^ waves. In pulmonary artery smooth muscle cells (PASMCs) it has been proposed that nicotinic acid adenine dinucleotide phosphate (NAADP) triggers increases in cytoplasmic Ca^2+^ via L-SR junctions, in a manner that requires initial Ca^2+^ release from lysosomes and subsequent Ca^2+^-induced Ca^2+^ release (CICR) via ryanodine receptor (RyR) subtype 3 on the SR membrane proximal to lysosomes. L-SR junction membrane separation has been estimated to be *<* 400 nm and thus beyond the resolution of light microscopy, which has restricted detailed investigations of the junctional coupling process. The present study utilizes standard and tomographic transmission electron microscopy to provide a thorough ultrastructural characterization of the L-SR junctions in PASMCs. We show that L-SR nanojunctions are prominent features within these cells and estimate that the junctional membrane separation and extension are about 15 nm and 300 nm, respectively. Furthermore, we develop a quantitative model of the L-SR junction using these measurements, prior kinetic and specific Ca^2+^ signal information as input data. Simulations of NAADP-dependent junctional Ca^2+^ transients demonstrate that the magnitude of these signals can breach the threshold for CICR via RyR3. By correlation analysis of live cell Ca^2+^ signals and simulated Ca^2+^ transients within L-SR junctions, we estimate that “trigger zones” with a 60–100 junctions are required to confer a signal of similar magnitude. This is compatible with the 130 lysosomes/cell estimated from our ultrastructural observations. Most importantly, our model shows that increasing the L-SR junctional width above 50 nm lowers the magnitude of junctional [Ca^2+^] such that there is a failure to breach the threshold for CICR via RyR3. L-SR junctions are therefore a pre-requisite for efficient Ca^2+^ signal coupling and may contribute to cellular function in health and disease.

## 1 Introduction

The importance of cytoplasmic nanojunctions to cellular signaling and thus to the modulation of cell function was recognised several decades ago [1, 2], henceforth the extent to which cellular nanospaces may contribute to the regulation of cell function received little attention. Nevertheless there is now a growing recognition of the widespread occurrence and functional significance of cytoplasmic nanospaces in cells from species across several kingdoms [3–9].

In this respect, membrane-membrane junctions between lysosomes and the sarcoplasmic reticulum (L-SR junctions) are of particular interest; not least because they have been hypothesized to couple calcium signaling between these two organelles [10, 11].

That L-SR junctions may play an important role in cellular Ca^2+^ signaling was uncovered through early studies on the Ca^2+^ mobilizing messenger nicotinic acid adenine dinucleotide phosphate (NAADP) [12], which demonstrated that NAADP released Ca^2+^ from a store other than the sarco/endoplasmic reticulum (S/ER) [13], that could then trigger further Ca^2+^ release from the S/ER by Ca^2+^-induced Ca^2+^ release (CICR) [14–17]. A major advance in our understanding was then provided by the demonstration that the NAADP-released Ca^2+^ was from an acidic lysosome-related store [10,18,19] in a manner that requires two pore segment channel subtype 2 (TPC2) [20]. However, studies on pulmonary arterial smooth muscle cells (PASMCs) had also identified a significant specialization, namely L-SR nanojunctions. It was hypothesized not only that these nanojunctions were necessary for coupling between lysosomes and the SR but that they could both coordinate and restrict their relationship to the SR by preferentially targeting ryanodine receptors while excluding inositol 1,4,5-trisphosphate (IP_3_) receptors [10, 11]. Importantly, NAADP-dependent Ca^2+^ bursts primarily arise from lysosomes in the perinuclear region of PASMCs and appear to promote rapid, local Ca^2+^ transients that are of sufficient size to activate clusters of SR resident ryanodine receptor subtype 3 (RyR3) and thus initiate, in an all-or-none manner, a propagating global Ca^2+^ wave [10, 11].

The specialization of the proposed L-SR junction is likely important in smooth muscle cell physiology, e.g., in vasomotion, and in the recycling of organelles and programmed cell death by autophagy [21]. However, L-SR junctions may also make as yet unforeseen contributions to vascular pathologies as highlighted by the fact that Niemann-Pick disease type C1 results, in part, from dysregulation of lysosomal Ca^2+^ handling [22] and is known to precipitate portal hypertension [23], while other lysosomal storage diseases (e.g., Pompe and Gaucher disease) accelerate pulmonary arterial hypertension [24, 25]. Moreover, observed hypertension is often associated with dysfunction of cholesterol trafficking [26], increased plasma cholesterol levels, vascular lesion formation, atherosclerosis/thrombosis and medial degradation [23, 27]. Therefore lysosomal Ca^2+^ signaling is of considerable clinical interest. That L-SR junctions may be of further significance to pathology is also evident, for example, from the fact that in the pulmonary artery smooth muscle L-SR junctions underpin Ca^2+^ waves initiated by endothelin 1, the levels of which are elevated in pulmonary hypertension, systemic hypertension and atherosclerosis [28, 29]. An understanding of how specific Ca^2+^ signals are functionally initiated therefore has important translational implications.

Lysosomal Ca^2+^ regulation has been of current interest in several recent research and review articles (e.g., [30–34]), yet the mechanism by which Ca^2+^ signals are generated by the endolysosomal system has not yet been modeled in a truly quantitative manner. Given the likely importance of L-SR junctions to Ca^2+^ signaling in health and disease, we sought to determine whether this nano-environment would indeed be able to effectively generate the previously observed NAADP-induced Ca^2+^ signals.

Due to the minute spatial scale of the nanojunctions generating the primary Ca^2+^ signals, accurate investigation of dynamic signaling within these spaces cannot be achieved with currently available instrumentation. To overcome this issue, we took an integrative approach by combining our own electron microscopy of L-SR nanojunctions, existing kinetic data on the Ca^2+^ transporters and buffers, and prior knowledge of the NAADP-induced Ca^2+^ signal features into a quantitative model of a typical L-SR nanojunction. This model is based on stochastic simulations of intracellular Ca^2+^ diffusion by Brownian motion implemented using the particle simulator MCell (freely available at mcell.org) [35–37]. In particular, we set out to verify the following hypotheses in PASMCs: (1) L-SR nanojunctions should be observable in the ultrastructure of these cells, (2) NAADP induces sufficient Ca^2+^ release from the lysosome to initiate activation of RyR3 embedded in the junctional SR, and (3) the combined effect of activation of L-SR nanojunctions in a cytoplasmic “trigger zone” determines the threshold of global [Ca^2+^]_i_ for the biphasic release process.

In the present manuscript we have verified the existence of L-SR nanojunctions within the ultrastructure of PASMCs, and shown that lysosomes can release sufficient Ca^2+^ to activate CICR via RyR3 clusters embedded in the junctional SR. Perhaps most importantly, we show that L-SR coupling is determined both by the integrity of L-SR junction on the nanoscale and the quantal summation of Ca^2+^ release from multiple, activated junctional complexes.

## 2 Results

### 2.1 NAADP-induced Ca^2+^ signals within isolated pulmonary artery smooth muscle cells

The relevant background findings that stimulated the development of the work presented here were first reported by Evans’ group in [17] and [10], and are summarized in figure 1.

**Figure 1.**
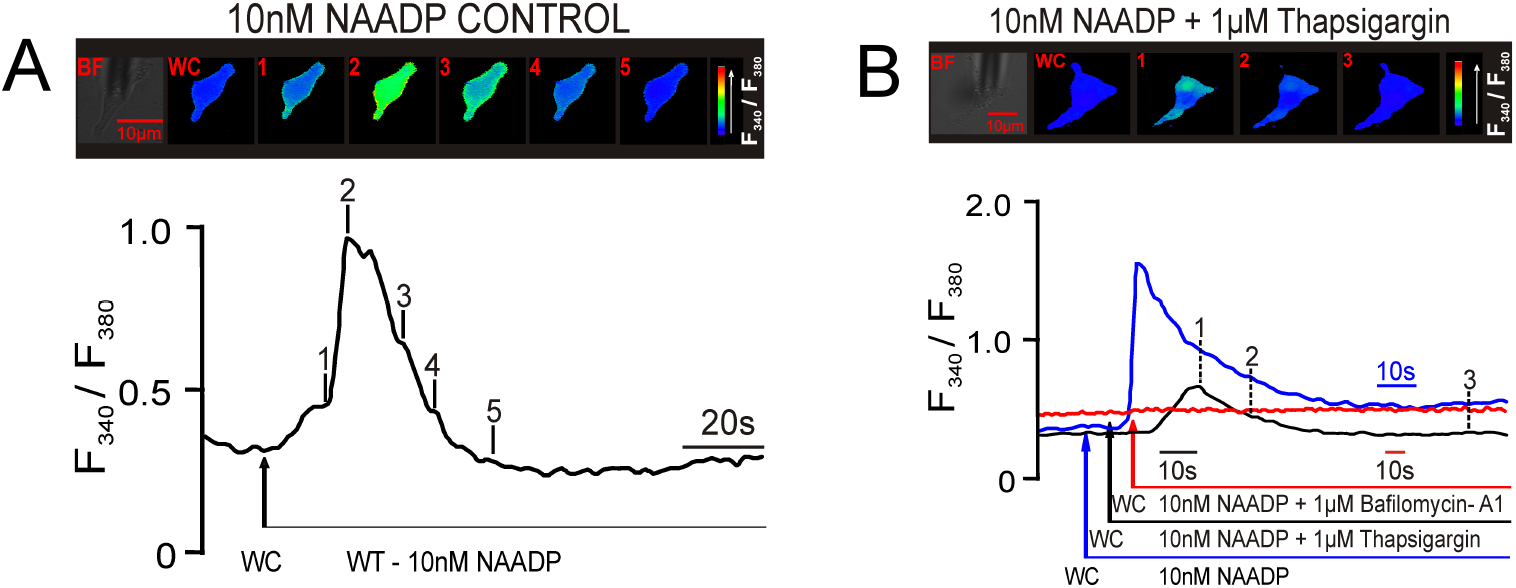
**A, B,** Upper panels show a series of pseudocolour images of the Fura-2 fluorescence ratio (F340/F380) recorded in two different pulmonary artery smooth muscle cell during intracellular dialysis of 10nM NAADP before (A) and after (B) depletion of SR stores by pre-incubation (30 min) with 1 *µ*M thapsigargin. Note the spatially localized ‘Ca^2+^ bursts’. A, Lower panel shows the record of the Fura-2 fluorescence ratio against time corresponding to the upper left panel of pseudocolours images; note the discrete shoulder in the rising phase of the F340/F380 ratio that corresponds to the initial ‘Ca^2+^ bursts’. B, Lower panel shows paired responses to 10 nM NAADP under control conditions (black), following depletion of SR Ca^2+^ stores with 1 *µ*M thapsigargin (blue) and following depletion of acidic stores with 1 *µ*M bafilomycin-A1 (red). Scale bars: 10 *µ*m.

The example record in figure 1A highlights the fact that NAADP appears to activate a two-phase Ca^2+^ signal, which can exhibit an identifiable “shoulder” during the initial rising phase of the signal (figure 1A, time point 1), followed by a second faster phase of signal amplification (figure 1A, time point 2). It is notable that the delay to the initiation of the second phase of amplification is variable [10, 11, 17] and due to this fact the shoulder is not always evident (see for example figure 1B, lower panel). Previous studies have demonstrated that this two-phase response results from initiation by NAADP of Ca^2+^ bursts from lysosome-related stores [10] in a manner that requires TPC2 [38] and that Ca^2+^ bursts are subsequently amplified by CICR from the SR via clusters of RyR3 [11,17]. Panel B of this figure illustrates the fact that prior depletion of SR stores by pre-incubation with thapsigargin (1 *µ*M) blocks the amplification phase, while depletion of acidic Ca^2+^ stores with bafilomycin abolishes the entire NAADP-induced Ca^2+^ signal. The previous studies cited above provided calibrated estimates of the changes in intracellular [Ca^2+^] input into the model below.

Based on earlier optical microscopy work, like the data in figure 1, and immunofluorescence results, it has been proposed that, for the lysosomal Ca^2+^ release to trigger CICR, L-SR nanojunctions are required and that they consist of apposing patches of lysosomal and SR membranes separated by a narrow space of nano-scale dimension [10, 17]. These studies led to an upper limit of 400 nm for the separation of the junctional membranes. We propose that L-SR junctions do indeed represent cellular nanojunctions and that they might play a role of accentuating Ca^2+^ gradients, akin to that of plasma membrane (PM)-SR junctions that are pivotal in the process of SR Ca^2+^ refilling during asynchronous [Ca^2+^] waves [39]. We furthermore hypothesize that in order for these nanojunctions to appropriately regulate Ca^2+^ signaling, they must be separated by a distance of approximately 20 nm and be of the order of a few hundred nm in lateral dimensions, as inferred from previous studies on PM-SR junctions [10, 11].

### 2.2 Ultrastructural characterization of L-SR nanojunctions

To identify lysosomes, SR regions and L-SR nanojunctions, we recorded and surveyed 74 electron micrographs of rat pulmonary arterial smooth muscle taken from samples prepared as described in the Materials and Methods section. The images in figure 2 provide a set of examples. Since we were aiming to detect L-SR junctions, namely close appositions of the lysosomal and SR membranes, immuno-gold labeling of lysosomes was prohibited, since this technique compromises membranes definition by electron microscopy to the extent that we would be unable to assess junctional architecture. Instead, in images like those in figure 2, lysosome identification was accomplished by relying on the knowledge of lysosomal ultrastructural features, which has accumulated over the past 50 years since the discovery of the lysosomes (see for example, [40]).

**Figure 2.**
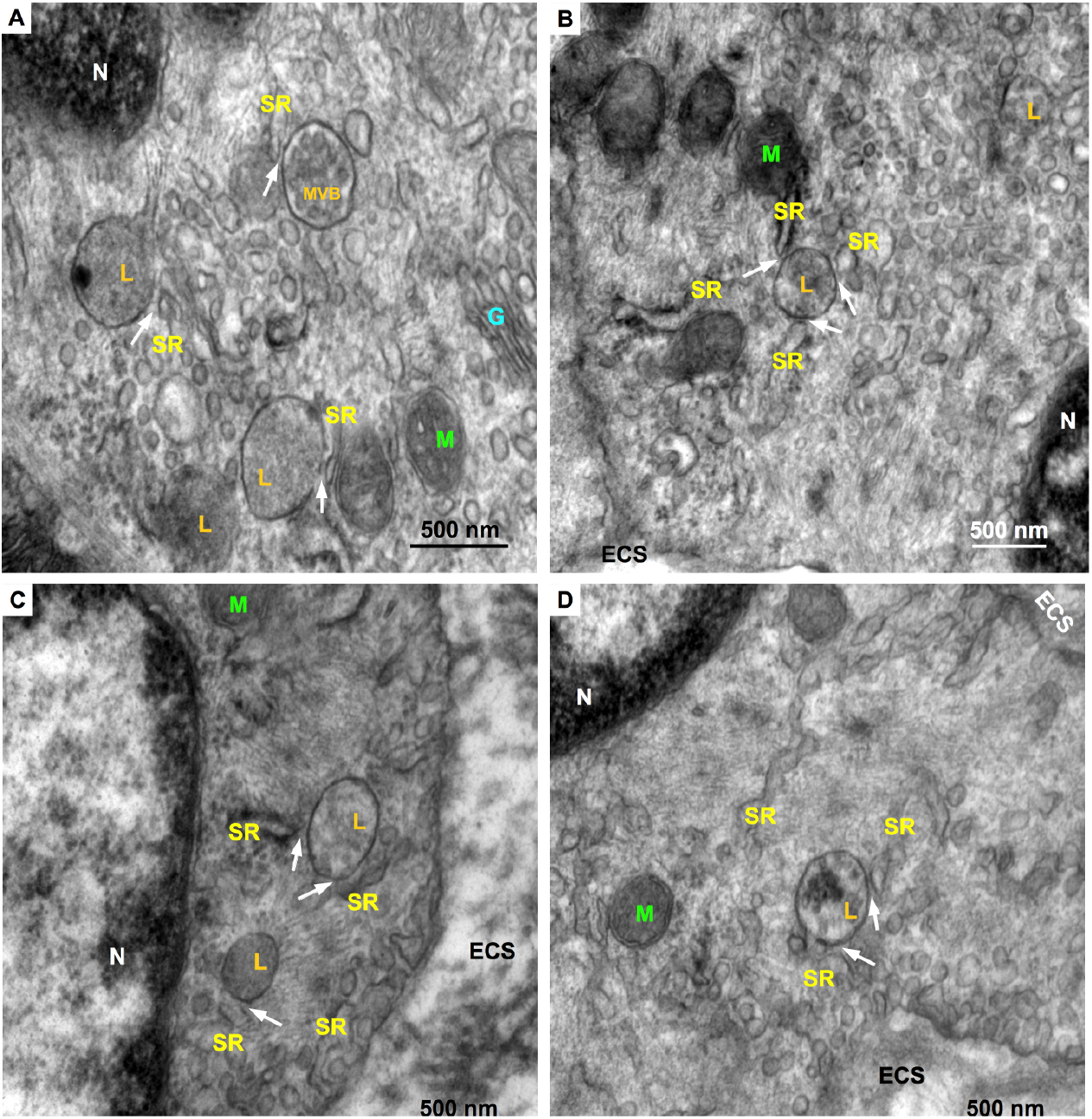
Representative electron micrographs of rat pulmonary artery SMC regions containing lysosomes (L), several SR cisterns, and including several examples of L-SR junctions (arrows). Also indicated are nuclei (N), Golgi apparatus (G), mitochondria (M), a multivesicular body (MVB) and extra-cellular space (ECS). Scale bars = 500 nm. Magnifications: A,C 80,000×, B, 60,000×, D, 70,000×.

In standard (2D) transmission electron microscopy (TEM) images, lysosomes typically appear as elliptical structures bound by a single lipid bilayer, a feature that distinguishes them from mitochondria. Depending on the lysosomal system stage, they also tend to have a more or less uniformly electron-dense interior as compared to the surrounding cytosol [41]. They can be distinguished from endosomes by their larger size and darker lumen and they differ from peroxisomes, since the latter usually display a geometrically distinct and markedly darker structure called “crystalloid” in their lumen. Moreover, it would appear that peroxisomes are found far more frequently in liver, kidney, bronchioles and odontoblasts than in other cell types (see, for example, [40,42,43]). Occasionally, organellar remnants are still visible inside these ovals, a characteristic that identifies them as multi-vesicular bodies (“MVB” in figure 2A). As it is at times questionable whether MVB’s are late endosomes or endosome-lysosome hybrids (compare, for example, [40] and [44]), we have excluded organelles (3 in total) with such characteristic from our statistical count.

From each of the relevant smooth muscle regions surveyed, we obtained high-resolution images of areas containing lysosomes and L-SR junctions in order to quantitatively characterize them (arrows in figures 2 and 3A). Using a software graphics editor (inkscape.org) and the image scale bar as a calibration gauge, we measured the lysosome size, as the length of the major and minor axes of their elliptical 2D projections (in orange and grey, respectively, in figure 3A), the L-SR widths, that is the distance between lysosomal and SR membranes at places where the two were about 30 nm or closer to each other (in purple in figure 3A), and the L-SR extensions as a percentage ratio between the junctional SR and the lysosomal membranes (in turquoise in figure 3A). From these measurements, we extrapolated the 3D junctional SR extension, both as a percentage of the lysosomal surface and as a length in nm. The histograms displayed in figure 3B–D visually summarize the data collected from the image analysis. The mean and standard deviation values of the measured parameters are reported in table 1.

**Figure 3.**
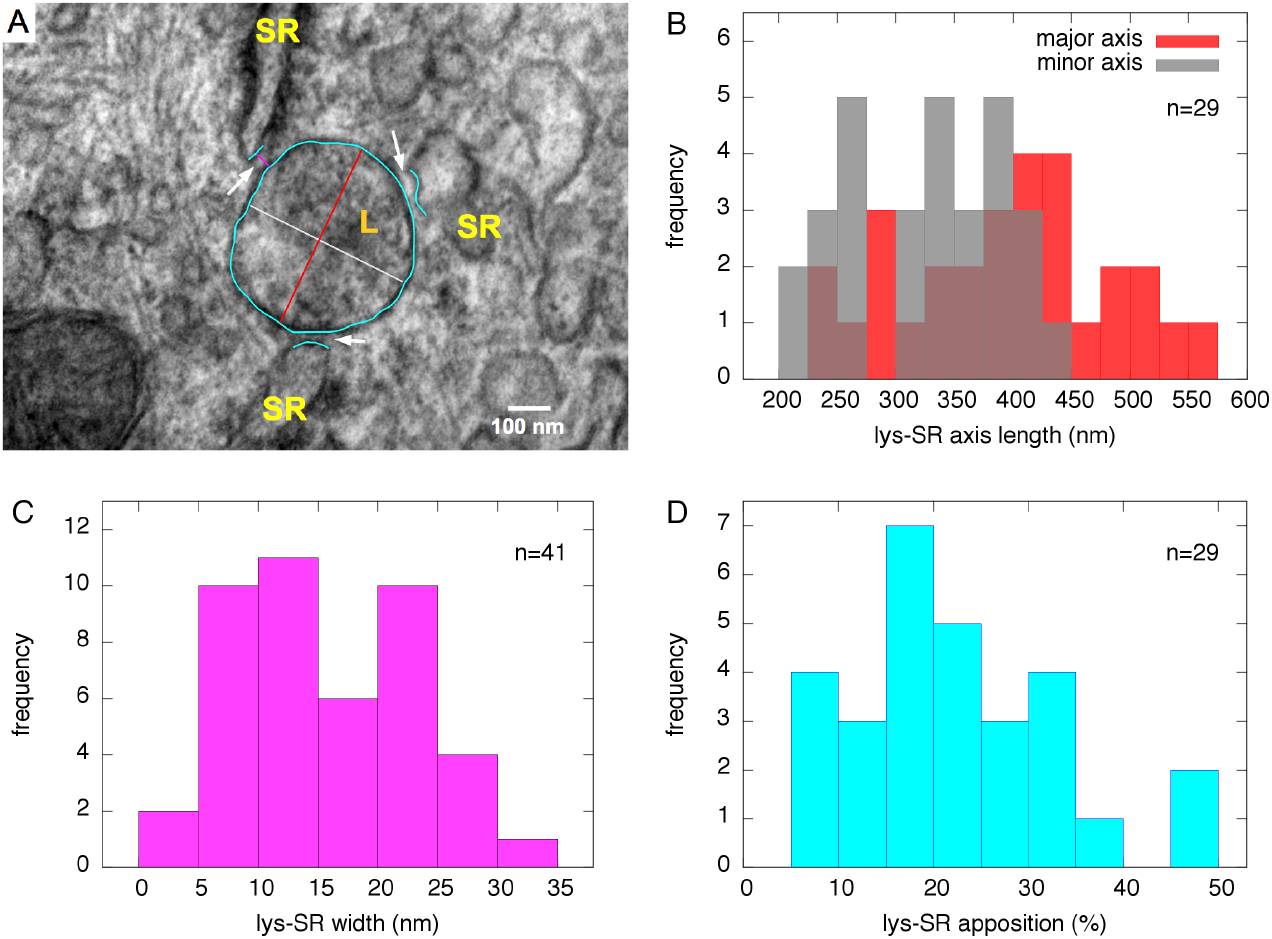
**A,** High magnification (150,000×) electron micrograph of a region of Figure 2B containing 3 L-SR junctions (arrows); coloured tracings as shown were used to measure lysosome dimensions, L-SR widths and extensions. Scale bar = 100 nm. **B–D,** Histograms showing distribution of several relevant lysosomal and L-SR junctional parameters, used to characterize the junctions and inform the quantitative model. B, lysosomal dimensions as major and minor axes of oval shape in micrographs; C, L-SR junctional width; D, percentage apposition between junctional SR and lysosome perimeter as projected in 2D micrographs.

**Table 1.** L-SR junction characterization parameters

| parameter | mean ± SD | notes |
| --- | --- | --- |
| lysosome minor axis | (325 ± 65) nm | $n = 29$ |
| lysosome major axis | (398 ± 91) nm | $n = 29$ |
| L-SR width | (16 ± 7) nm | $n = 41$ |
| L-SR overlap | (23 ± 13) % | $n = 29$ |
| L-SR lateral extension | (262 ± 209) nm | calculated from data above |
| 3D L-SR overlap | (13 ± 15) % | extrapolated from 2D measurements |

Mean and standard deviation values of L-SR junction parameters, calculated from data as in Figure 3B–D.

Estimates of the various parameters gathered in this phase of the study were used as a basis to build a 3D software object to represent a typical L-SR junction. This reconstruction was then used to design the simulations mimicking Ca^2+^ diffusion in the L-SR nanojunctions, as is described below.

#### 2.2.1 **Tomography**

To gather more direct information on the 3D morphology of L-SR junctions, we acquired a set of tomograms of those regions from the same sample blocks used to obtain the images in figure 2. In figure 4, we report snapshots from one of the tomograms; in these stills, we have also traced out parts of one lysosome and the closely apposed SR region that together form a L-SR junction. These tomograms are very helpful in clarifying the detailed morphology of L-SR junctions and informing on the spatial variability of the SR network. For example, while the SR segment shown in a single 2D tomographic scan (figure 4A) would appear to be continuous and part of a large SR compartment, it actually branches out into narrower cisterns as revealed by 3D tomographic reconstruction (figure 4B). Furthermore, it is interesting to observe the fact that one extension of the SR appears to couple with multiple organelles. Thus, the 3D views generated by tomography are paramount for demonstrating the presence of a true junctional complex and for the design of a prototypical L-SR environment as a software mesh object, on which we may simulate the NAADP-mediated localized Ca^2+^ release.

**Figure 4.**
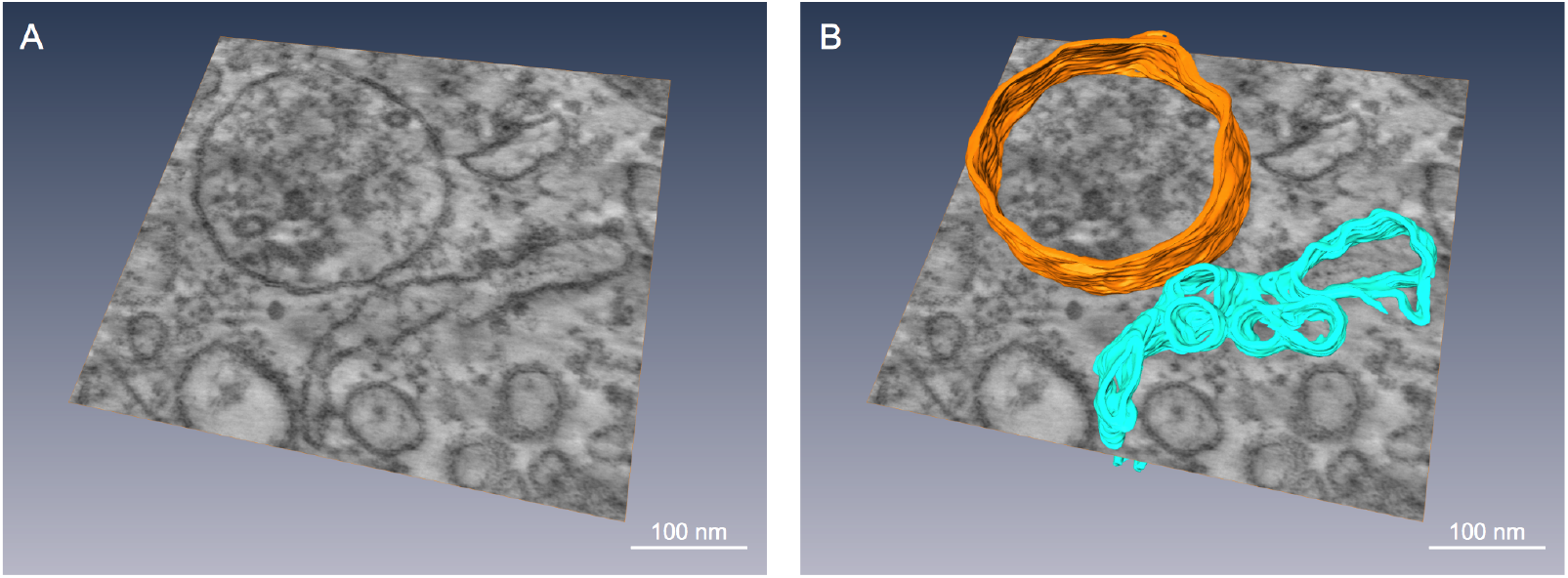
**A,** Snapshot from a TEM tomogram of a L-SR region of rat pulmonary artery smooth muscle, illustrating, among other things, a single SR extension apparently forming junctions with several lysosomes. Magnification = 62, 000×. **B,** Same snapshot shown in A, but with a lysosome (orange) and a portion of SR (turquoise) partially traced out in 3D. These pseudo-colour tracings underscore how the SR can appear as a large cistern in a given plane, but can actually branch out in different directions when viewed in 3D. Scale bars *≈* 100 nm.

### 2.3 **Quantitative model**

The model aims to verify whether NAADP-induced Ca^2+^ release from the lysosomal system could be responsible for the localized Ca^2+^ signal preceding the global Ca^2+^ wave (see figure 1 and [10]), which triggers a propagating wave by CICR via RyR3s localized at L-SR junctions, as predicted by earlier observations [10].

#### 2.3.1 Generation of the signal’s “shoulder” by lysosomal Ca^2+^ bursts

To understand the generation of the signal “shoulder”, such as that observed at time point 1 in the lower panel of figure 1A, by Ca^2+^ bursts within L-SR nanojunctions, let us note that its magnitude corresponds to the difference in [Ca^2+^] between the resting level of ≈ 100 nM [17] prior to NAADP stimulation (up to the point, at which NAADP enters the cytoplasm under the whole-cell configuration (WC) in figure 1) and the value of ≈ 400 nM [17], corresponding to [Ca^2+^]_i_ at time point 1 in the example record shown in figure 1A, lower panel (these concentration values were obtained via a standard calibration procedure [17]). This leads to a Δ[Ca^2+^]_shoulder_ of approximately 300 nM [17]. Let us now estimate the number of lysosomes required to generate a Δ[Ca^2+^]_shoulder_ of such magnitude in a PASMC, given the following assumptions:

1. That the luminal [Ca^2+^] of a lysosome, [Ca^2+^]_lys_, is in the range of 400–600 *µ*M, as determined in mouse macrophages [45] and that it is homogeneous across the lysosomal population;
2. That the Ca^2+^ release rate during bursts is ≈ 10^6^ ions/s (based on findings in [46], but see next section for a detailed analysis on the rate time-variation) and that it becomes negligible for values of [Ca^2+^]_lys_ below ≈ 80 *µ*M, as suggested by the single channel kinetics of TPC2 signaling complexes in lipid bilayer studies [46];
3. That lysosomes are spheres with a radius of ≈ 180 nm, as gathered from our EM characterization (see figures 2, 3, 4, and table 1), and hence that their volume is *V*_lys_ = (4*π/*3)(1.8 × 10*^−^*^7^ m)^3^ = 2.4 × 10*^−^*^17^ L;
4. That a smooth muscle cell cytosolic volume can be calculated as *V*_cyt_ = 2.4 × 10*^−^*^12^ L by modeling a cell as a 130-*µ*m-long cylinder, 6 *µ*m in diameter [47], and accounting for nuclear, SR, mitochondrial and lysosomal volumes.

With these assumptions accepted, from points 1. and 2. above we gather that a lysosome can release a potential Δ[Ca^2+^]_lys_ = (400 to 600)*µ*M − 80*µ*M = 3.2 × 10*^−^*^4^ M to 5.2 × 10*^−^*^4^ M into the cytosol (as we elaborate in the next section, this should take ≈ 0.03 s, a time frame that ensures that released Ca^2+^ can be considered unbuffered). Each lysosome contribution to [Ca^2+^]_i_, Δ[Ca^2+^]_i,lys_, can then simply be calculated by taking the lysosome-to-cell volume ratio into account:

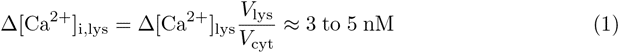

Therefore, we can calculate the number of lysosomes that may contribute to the magnitude of Δ[Ca^2+^] shoulder as

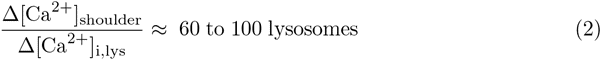

In summary, between 60 and 100 lysosomes would be necessary (and possibly sufficient) to provide a Δ[Ca^2+^]_shoulder_ of 300 nM, which is typically observed during the localized Ca^2+^ release phase of the NAADP-induced Ca^2+^ signals.

How many lysosomes do we actually expect to be in a PASM cell of our sample tissue? We can obtain a rough estimate of this number from the TEM imaging we performed for this study. In each 80-nm-thick TEM sample section, we see 5–10 lysosomes/cell. Lysosomes are predominantly localized to the perinuclear region of the cytoplasm, but seldom in the subplasmalemmal area, consistent with previous observations by optical microscopy [10, 11]. If we simplify the geometry of a typical smooth muscle cell to a 130-*µ*m-long cylinder with radius of 6 *µ*m, and considering that lysosome radii are around 180 nm, as mentioned above, it is reasonable to assume that separate sets of lysosomes would be observable in TEM images taken at distances into the sample of about 180 nm from each other. Neglecting the slices within one lysosomal diameter of the cylinder surface—given the near total lack of observed lysosomes in those subvolumes—then our images suggest that we can expect a total of about 130 lysosomes/cell. It is encouraging that we obtain from this count a higher number than the 60–100 lysosome range we derived in equation (2), in that on the one hand it is plausible to think that not all of the lysosomes in a cell may be activated in synchrony, nor may they all be involved in NAADP-mediated signaling, and on the other hand experience tells us that evolution has built in some redundancy of function in order to provide a threshold and also a margin of safety for the generation of this type of Ca^2+^ signals. Moreover, the estimated number of junctions is based on a value for [Ca^2+^]_lys_ determined in macrophages, and it is plausible that the total releasable [Ca^2+^]_lys_ in a PASMC may differ from that value and that it may also vary over the lysosome population. Lastly, the uncertainty in the number of lysosomes/cell evidenced from the electron micrographs as described above may also contribute to this discrepancy.

#### 2.3.2 L-SR junctional Ca^2+^ signal

In the hypothesized model outlined in [10, 11], Ca^2+^ bursts activate SR resident RyR3 channels within L-SR junctions and thus initiate a propagating Ca^2+^ wave by CICR. The stochastic simulations developed here attempt to reproduce the phenomenon of the generation of [Ca^2+^] transients within individual L-SR junctions, considering the Ca^2+^ release kinetic requirements for lysosome-resident TPC2 signaling complexes and the rate of Ca^2+^ capture by the SERCA2a localized on the neighbouring SR membrane [48], and to link these junctional transients to the observed bursts.

The thorough quantitative image analysis described in the previous section yields critical information for our first modeling phase, in which we built a dimensionally accurate virtual lysosome and a portion of the SR system, closely apposing the lysosome (figure 5), so as to reproduce a NAADP-triggered Ca^2+^ signal within the nanospace of a representative L-SR junction as faithfully as possible.

**Figure 5.**
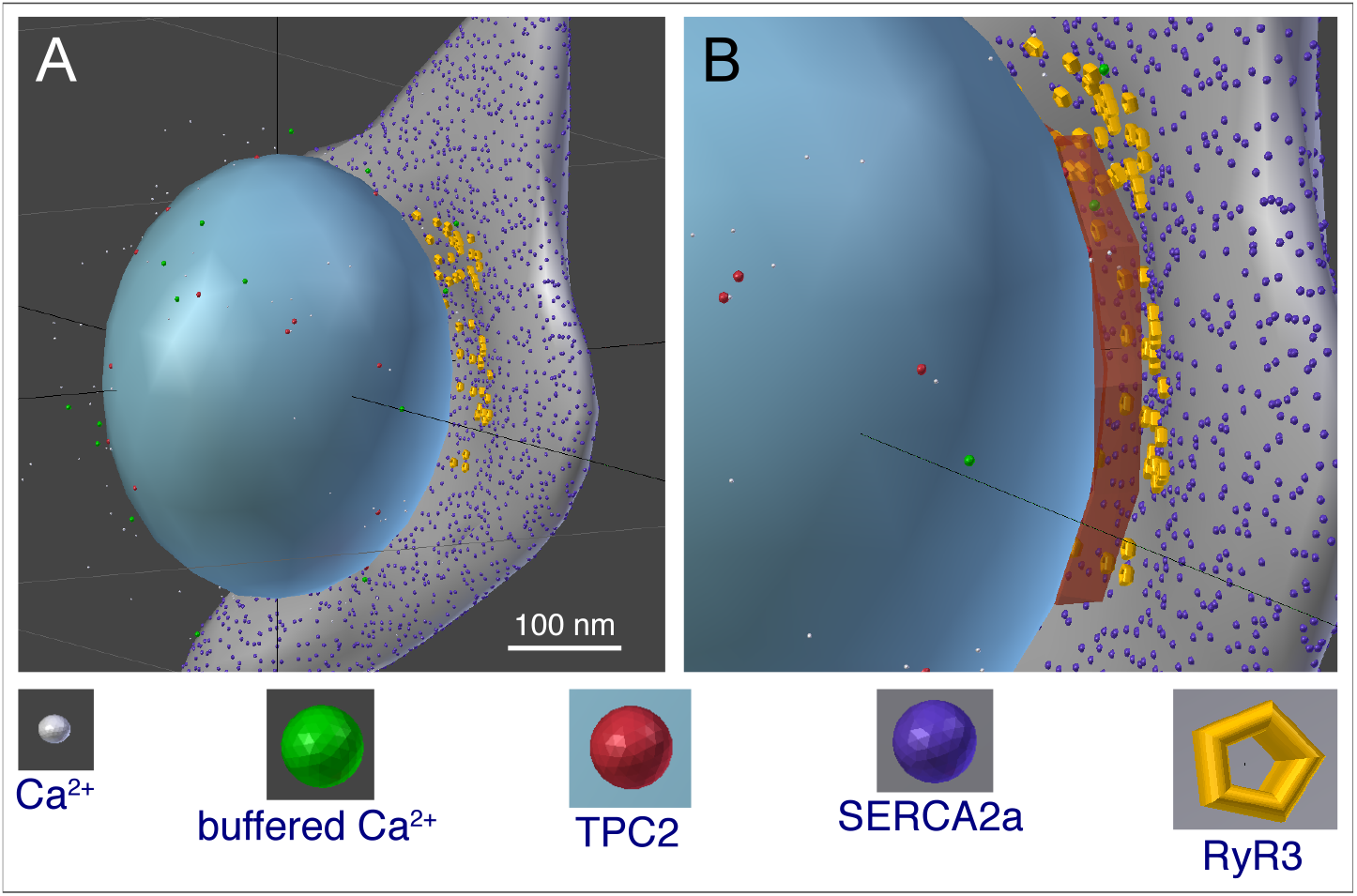
**A and B**, 3D software reproduction of a lysosome closely apposed to a portion of SR, thereby forming an *≈* 20-nm-wide L-SR nanojunction; this rendering was inspired by a series of observations from micrographs as in Figure 2 (grey object = SR, blue object = lysosome). Included are relevant molecules traversing Brownian motion trajectories produced by the model simulations (see symbol legend below the panels). B, enlarged view of the L-SR junctional region, in which we have displayed the volume object (rust-coloured box) used to measure the [Ca^2+^]_NJ_ transients like the ones reported in figure 7. Scale bar *≈* 100 nm. The model geometry and code files are available from the corresponding author.

From the available literature we estimated the number of TPC2 and SERCA2a likely distributed on the lysosome and SR membranes, respectively. We obtained the former number by dividing the macroscopic whole-lysosome conductance, calculated from the current values reported in a recent study on TPC2-mediated Ca^2+^ current in isolated lysosomes [49], by the single channel conductance determined in [46], thus estimating that a typical lysosomal membrane may contain *≈* 20 TPC2. To obtain this value, we used the experimental condition data provided in [49] to extract values for the ionic potential, *E*_ion_, across the lysosomal membrane. We then employed the authors’ current-voltage (*I*-*V*_m_) data to compute whole-lysosome conductance values (*g*_WL_) as a function of the applied membrane potential, *V*_m_, from Ohm’s law: *I* = *g*_WL_(*V*_m_ − *E*_ion_).

Moreover, we have estimated the density of SERCA2a on the SR within L-SR junctions to be equivalent to that previously predicted for PM-SR junctions as approximately 6250/*µ*m^2^ [39]. Other input data for the model are the estimates of lysosomal volume and of the [Ca^2+^]_lys_, from which we calculated the actual number of ions in the lysosome prior to the beginning of Ca^2+^ release (see previous section). This, in turn, was used in the extrapolation of the TPC2 complex Ca^2+^ release rate as a function of time, as follows.

A recent electrophysiological study of the Ca^2+^ conductance of TPC2 signaling complex provides valuable information regarding its biophysical properties and, importantly, highlights the fact that, for a given relatively low activating concentration of NAADP (10 nM, as in figure 1), the channel open probability appears to depend on [Ca^2+^]_lys_ (this is likely due to a partial neutralization of the electrochemical potential across the lysosomal membrane) [46]. We used the Ca^2+^ conductivity measured in [46] and values of the lysosome membrane potential ([50]) to calculate the maximal Ca^2+^ current, *I*_max_, as 2.4 × 10^6^ ions/s. We then applied a weighted quadratic fit to the channel’s open probability (*P*_o_) as a function of the [Ca^2+^]_lys_ data in [46] with constraints that the *P*_o_ would tend to zero at [Ca^2+^]_lys_ = 80 *µ*M, based on the observation that below [Ca^2+^]_lys_ = 100 *µ*M essentially no single channel openings were observed [46]. Another constraint for the fit was that the curve be within the standard deviation value of *P*_o_ at the highest reported [Ca^2+^]_lys_ = 1 mM. From the quadratic fit, we then obtained our own *P*_o_-vs-[Ca^2+^]_lys_ table (plotted in figure 6A) and assumed that at the beginning of Ca^2+^ release, the Ca^2+^ current would be *P*_o,max_ *× I*_max_ at the maximal luminal concentration until the luminal concentration decreased to the next point in the *P*_o_-vs-[Ca^2+^]_lys_ relationship. At this point, the release rate decreases to a new (lower) *P*_o_ × *I*_max_ until the luminal concentration reaches the next lower point in the table, and so on until *P*_o_ = 0 at [Ca^2+^]_lys_ = 80 *µ*M. The Ca^2+^ release rate as a function of time obtained in this manner is shown in figure 6B.

**Figure 6.**
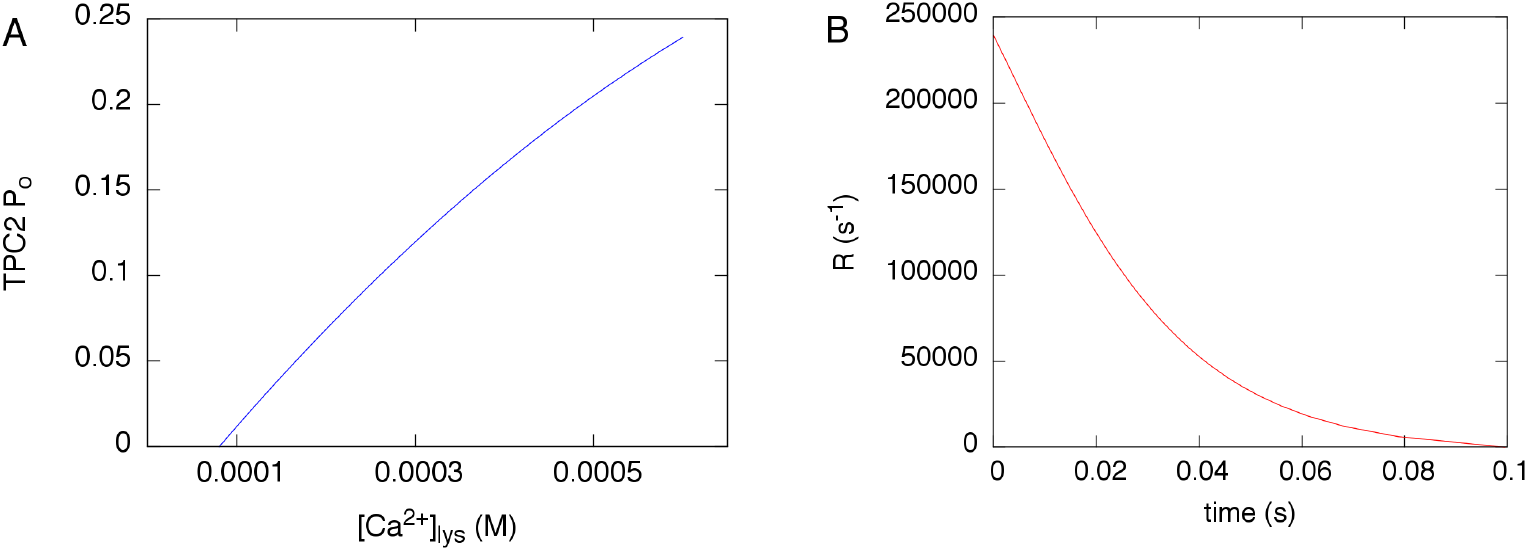
**A,** open probability *P*_o_ of the TPC2-dependent Ca^2+^ conductance reproduced from a quadratic fit to the data reported in [46]. **B,** Ca^2+^ release rate calculated as explained in the text, based on the *P*_o_ in A.

To implement an approximated SERCA pump action, we used a simplified version of a multi-state kinetic model developed in [51], which was further informed by more recent isoform-specific studies [52, 53]. In particular, since we are interested in simulating the SERCA2a Ca^2+^ uptake only in terms of its influence on the shaping of the junctional [Ca^2+^] transient, we have opted not to implement the steps of the multistate model that deal with the Ca^2+^ unbinding from the SERCA on the SR luminal side of the pump. In brief, the reactions SERCA2a undergo are: (1) binding/unbinding of the first Ca^2+^; (2) binding/unbinding of the second Ca^2+^.

The Ca^2+^ diffusivity was obtained from studies, in which it was concluded that given the known kinetics of typical Ca^2+^ buffers, the range of free Ca^2+^ after it enters a cell can be up to 200 nm. Therefore, due to the nano-scale of our system, the trajectories of Ca^2+^ released by TPC2 signaling complexes in the simulations are governed by the measured diffusivity of free Ca^2+^(2.23 × 10*^−^*^10^ m^2^/s; [54, 55]). Once Ca^2+^ are buffered they acquire the measured diffusivity of the buffers (8.4 × 10*^−^*^11^ m^2^/s; [56]).

We summarize the quantitative model input data in table 2.

**Table 2.** Quantitative model input data

| quantity | value | source |
| --- | --- | --- |
| $\Delta[\text{Ca}^{2+}]_{\text{shoulder}}$ | 300 nM | [17] and this article |
| $[\text{Ca}^{2+}]_{\text{lys}}$ | 400–600 $\mu\text{M}$ | [45] |
| lysosomal average radius | 180 nm | this article, figure 3 |
| (cylindrical) SMC radius | 6 $\mu\text{m}$ | [47] |
| SMC length | 130 $\mu\text{m}$ | [47] |
| SMC cytosolic volume | $2.4 \times 10^{-12}$ L | this article |
| TPC2 complexes per lysosome | 20 | [46, 49] and this article |
| TPC2 complex $\text{Ca}^{2+}$ release rate | variable | this article, figure 5 |
| SERCA2a surface density | $6250/\mu\text{m}^2$ | [39] |
| SERCA2a first $\text{Ca}^{2+}$ binding (unbinding) rate | $5 \times 10^8 \text{ M}^{-1}\text{s}^{-1}$ ($60 \text{ s}^{-1}$) | [51–53] |
| SERCA2a second $\text{Ca}^{2+}$ binding (unbinding) rate | $4 \times 10^8 \text{ M}^{-1}\text{s}^{-1}$ ($60 \text{ s}^{-1}$) | [51–53] |
| free $\text{Ca}^{2+}$ diffusivity, $D_{\text{free}}$ | $2.23 \times 10^{-10} \text{ m}^2/\text{s}$ | [54, 55] |
| $\text{Ca}^{2+}$ buffer diffusivity, $D_{\text{buffer}}$ | $8.4 \times 10^{-11} \text{ m}^2/\text{s}$ | [56] |

Summary of input parameters used in various phases of the quantitative model, and references to the origin of their values.

Using MCell as a stochastic particle simulator, we ran a number of simulations to represent the NAADP-mediated Ca^2+^ release that is supposed to occur at L-SR junctions in PASM experiments. Released Ca^2+^ is assumed to undergo Brownian motion in the surrounding space, including the L-SR nanospace. SERCA2a placed on the neighbouring SR surface may capture Ca^2+^ according to our approximation of their known multistate model. To determine how this regenerated cellular environment can shape a Ca^2+^ transient we “measured” the junctional [Ca^2+^] by counting the ions within the L-SR volume at any given time and dividing the number by the volume. The snapshots in figure 5 are part of the visual output of this phase of the work. The data to determine the simulated Ca^2+^ transients in the L-SR junctions were collected in a measuring volume placed between the lysosomal and SR membranes in the virtual L-SR junction (rust-coloured box in figure 5B).

Previous experimental findings about VSMC PM-SR junctions indicate that disruption of nanojunctions can have profound consequences on Ca^2+^ signaling properties of the cells. For example, when calyculin A was used to separate superficial SR portions from the plasmalemma of the rabbit inferior vena cava, it was observed that [Ca^2+^]_i_ oscillations would cease [57]. This was later corroborated by a quantitative model of the PM-SR junction’s role in the refilling of SR Ca^2+^ during oscillations [39]. In another study by the van Breemen laboratory, it was observed that the mitochondria-SR junctions of airway SMC displayed a variable average width as a function of the state of rest or activation of the cell [58]. It makes sense then to study the effect of changes in the junctional geometry on the transients generated by our simulations. Therefore, we ran several simulations, in which the separation between the lysosomal and SR membranes was increased from 10 nm to 100 nm in steps of 10 nm. In figure 7A, we report three sample transients from this set of simulations obtained using three different junctional membrane separations, as indicated in the inset legend. The value of each of the points graphed in this plot is the average value of 100 simulations, in each of which the random number generator within MCell is initiated with a different seed (see Materials and Methods). We report representative error bars as 3× the standard error, to convey the *>* 99% confidence interval of the data. As one would expect, the transient nanojunctional (NJ) Ca^2+^ concentration, [Ca^2+^]_NJ_, decreases in magnitude, as the junctional L-SR membrane-membrane separation increases, simply because the released Ca^2+^ has a larger junctional volume available over which to spread. However, it is important to make a quantitative comparison between this change in [Ca^2+^]_NJ_ and the [Ca^2+^]_i_ requirements to activate the putative RyR3 population of the junctional SR. This can be attained by analyzing the time scale of both the recorded and simulated Ca^2+^ signals.

**Figure 7.**
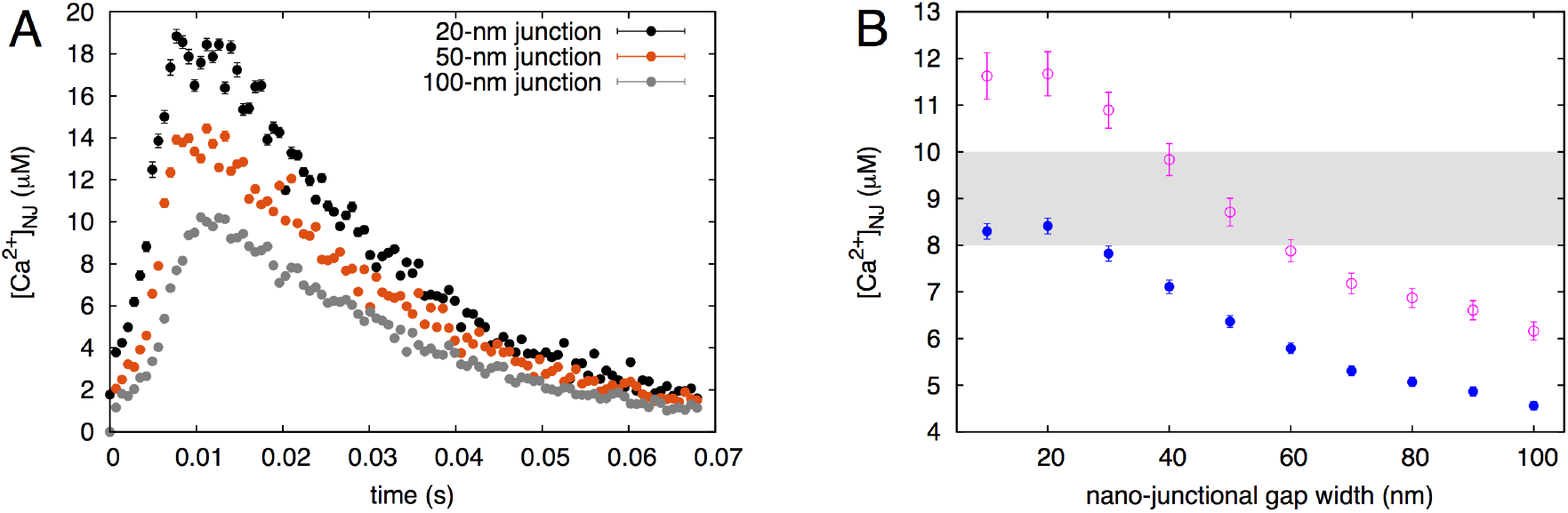
**A,** calculated nanojunctional [Ca^2+^] transient, [Ca^2+^]_NJ_, “measured” inside the volume of the recreated L-SR nanojunction shown in figure 5. To show the effect of changes in the junctional geometry, we report three transients calculated using different junctional widths of 20, 50 and 100 nm. **B,** [Ca^2+^]_NJ_ *vs* width of junction, concentration values are temporal averages of the transients as in panel A, calculated over an interval of 0.065 s (solid circles) and 0.038 s (empty circles); see text for explanation. The shaded area indicates the approximate threshold values for CICR at RyR3s [59].

#### 2.3.3 Reconciling the temporal scales in simulation and experimental results

In the final step of the development of our model, we analyzed the relationship between the time scale of the [Ca^2+^]_NJ_ transients resulting from our model (figure 7) and that of the observed Ca^2+^ signal in figure 1.

Let us note that the typical duration of the simulated transients, *t*_transient_, is about 0.06 s and recall that these represent Δ[Ca^2+^] above the resting [Ca^2+^]. On the other hand, the build up to the maximum value of Δ[Ca^2+^]_shoulder_ takes about 5 s (we refer to this time as *t*_shoulder_; figure 1). This interval is about two orders of magnitude larger than the duration of the simulated individual transients (figure 7A). One way to reconcile the hypothesis that L-SR junctions are at the base of the observed NAADP-induced Ca^2+^ signals such as the ones in figure 1 and in particular that the signal shoulder emerges from lysosomal Ca^2+^ release at L-SR junctions, is to bring forward the assumption that the signal shoulder may be the result of a sequential summation effect over many L-SR junctions, each working according to an all-or-none mechanism of Ca^2+^ release, and that the “firing” of one junction causes a cascading effect across the set of junctions that produce the shoulder. Then the duration of the shoulder upstroke can be expressed as:

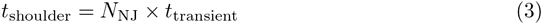

where *N*_NJ_ is the number of L-SR nanojunctions that yield Δ[Ca^2+^]_shoulder_. Since we calculated above that Ca^2+^ release from *N*_NJ_ ≈ 60–100 lysosomes would be necessary to produce the observed signal shoulder magnitude and have shown in the previous section that such release would need to take place at L-SR junctions, we can estimate *t*_transient_, the duration the [Ca^2+^]_NJ_ transient, by reversing equation (3) and obtaining *t*_transient_ = 0.05–0.08 s. It is noteworthy that this range of values is obtained in a manner completely independent of our simulation results, which yielded a similar range of values.

To gain quantitative insight into the effects of varying the junctional width, we then calculated the temporal average of the [Ca^2+^]_NJ_ over *t*_transient_ (using the middle value of the range calculated via equation (3)) and graphed it as a function of the junctional width. The result of this analysis is reported in figure 7B (solid blue circles). As we have anticipated at the end of the previous section, the decrease in magnitude of these data is to be expected, however in this plot we also indicated the range of [Ca^2+^]_i_ values (shaded area) over which maximum SR Ca^2+^ release via RyR3 is reported to occur in skeletal muscle [59]. This comparison underscores the important constraint played by the width of the L-SR junctions and indicates that, unless the membrane separation is kept below about 30 nm, it is not possible for [Ca^2+^]_NJ_ to breach the threshold for RyR3 Ca^2+^ release. Let us also point out that the [Ca^2+^]_NJ_ data in figure 7B would shift upward, toward concentration values that would make the junction more prone to promote RyR3 release, if the temporal average were taken over a shorter transient time, 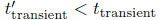 around the [Ca^2+^]_NJ_ peak. However, in that case equation (3) indicates that a larger *N*_NJ_ (than 80, picked as the middle of the 60–100 range) would have to contribute to the signal summation that results in a 5-second *t*_shoulder_. Interestingly, this possibility agrees with our lysosome count from TEM images (130 lysosomes/cell) and with the argument of natural redundancy we contemplated to explain the discrepancy between the calculated and observed lysosome number estimates. As an exercise we have recalculated the [Ca^2+^]_NJ_ time-averaged over 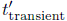 obtained using 130 lysosomes in equation (3) (empty purple circles in figure 7B), and this indeed shows that RyR3 Ca^2+^ release threshold would be cleared more readily. These observations suggest that activation of RyR3s in the L-SR junctions not only depends on the concentration of Ca^2+^ near them, but also on their exposure time to this concentration.

## 3 Discussion

### 3.1 Inter-organellar nanospaces

We have recently introduced the concept of the “pan-junctional SR”, which states that Ca^2+^ release and uptake at a family of specific nanojunctions connected by a continuous but variable SR lumen integrates cellular control over multiple functions [4]. The lysosome-sarco/endoplasmic reticulum (L-S/ER) junction is the most current junction to be considered in this context and exhibits perhaps the highest degree of plasticity of the family of nano junctions of the SR; the mechanism and function of lysosomal Ca^2+^ signaling is currently hotly debated [60].

By means of a thorough ultrastructural study in rat pulmonary artery smooth muscle, we have observed and characterized L-SR nano-junctions, which had been previously hypothesized on the basis of optical measurements of Ca^2+^ signals and optical immunocytochemistry [10, 11]. Our observations corroborate the previously reported finding (in [10] and [17]) that lysosomes in PASMCs tend to cluster in the perinuclear region, as is evident in our micrographs (e.g., figure 2A). We find that L-SR junctions are on average 15 nm in width (equivalent to our preliminary reports [8, 61] and to recent observations in cultured fibroblasts [33]) and extend approximately 300 nm in lateral dimensions, thereby involving about 15% of the lysosomal membrane (table 1).

### 3.2 Mechanism of NAADP Ca^2+^ signaling

In an effort to achieve quantitative understanding of the phenomenon of NAADP-mediated Ca^2+^ transients and verify the proposal that these may be generated in L-SR junctions [10, 17], we focused on one of the prominent features of these Ca^2+^ signals, namely the localized Ca^2+^ bursts that precede the propagating Ca^2+^ wave (figure 1), which we refer to as the signal shoulder (Δ[Ca^2+^]_shoulder_). In the first instance, we have estimated the potential contribution of local bursts of Ca^2+^ release from individual lysosomes to the elevation of global [Ca^2+^]_i_ observed during this shoulder in the experimental records. From this, and using the dimensions of a typical smooth muscle cell, we calculated that 60–100 lysosomes would be required to cause an elevation of comparable magnitude to the signal shoulder. This is lower than the estimate for the total number of lysosomes per cell, 130, we obtained from our ultrastructural study, but comparable in order of magnitude. We have already mentioned above a number of factors in favour of observing a greater number of lysosomes/cell than the estimated number required to generate the signal shoulder. In addition, several other elements contribute a degree of uncertainty to those estimates, such that their discrepancy may not be as large. We need to consider that *P*_o_ data for the Ca^2+^ conductance associated with the TPC2 signaling complex published in [46], on which we based our TPC2 rate table, show some variability according to the standard deviation bars, which, in turn, implies an uncertainty in the interpolated *P*_o_ (figure 5A). Moreover, we cannot exclude the possibility that these data reflect a contribution from multiple channels (*N P*_o_) rather than a purely single channel *P*_o_. Lastly, the standard deviation of the simulated transient (only the standard error is shown in figure 7A) and the variability in the experimental determination of the [Ca^2+^] sensitivity of RyR may allow for some uncertainty in the estimated number of nanojunctions.

To take this study further and understand whether the observed L-SR junctions could give rise to [Ca^2+^]_i_ transients of appropriate magnitude to trigger Ca^2+^ release from RyR3 channels on the junctional SR, we developed a quantitative stochastic model of Ca^2+^ dynamics in the junctional nanospaces. We have previously published a simplistic version of this model, which nonetheless captured the essential features of the problem and yielded an indication that such Ca^2+^ transients in the L-SR junctions would be possible [4]. However, the simulated transients we obtained displayed unphysiological features, such as an abrupt onset and decay. We show here that this was largely due to lack of a faithful representation of the open probability for the Ca^2+^ conductance of the TPC2 signaling complex. We have now combined experimental information on the biophysical properties of conductance and open probability (from [46]) and on the luminal [Ca^2+^] of the lysosomes (from [45]) to implement a more realistic Ca^2+^ release rate model in the simulations. As a consequence, we are able to output a physiologically meaningful junctional transient profile (figure 7A), and observe that the [Ca^2+^]_NJ_ transients generated within our model junctions—in turn, based on those observed in our TEM images—reach peaks of about 20 *µ*M. While we are still not able to measure these transients in individual L-SR junctions (peri-lysosomal Ca^2+^ probes are only recently becoming available—see for example [62]—and so far have not been used in smooth muscle cells, vascular or otherwise), it is worth noticing that the values we find are comparable to those recently measured in so-called Ca^2+^ hot spots in the mitochondria-ER junctions of neonatal ventricular cardiomyocytes (rat culture) [63] and in RBL-2H3 and H9c2 cells earlier [64]. These results therefore suggest that the hypothesis presented for the role of L-SR junctions in cellular Ca^2+^ signaling is certainly plausible.

### 3.3 Lysosomal trigger zone

Although our simulations were successful, comparison of the Ca^2+^ transients in figures 1 and 7 not surprisingly reveals a striking difference between the simulated Ca^2+^ kinetics of a single L-SR junction and those of the whole cell. This illustrates important aspects related to the concept of lysosome-SR “trigger zone” previously introduced by one of us (AME, [10]) and recently proposed as a facilitating factor in lysosomal-ER signaling in reverse, whereby Ca^2+^ release from ER compartments enables NAADP-mediated activation of acidic organelles [32].

Briefly, this concept suggests that in PSMCs clusters of lysosomes and closely apposed SR regions containing sets of RyRs may act together to form specialized intracellular compartments that transform NAADP-stimulated localized lysosomal Ca^2+^ release into cell-wide Ca^2+^ signals via RyR-supported CICR. If the firing of one junction were sufficient for subsequent initiation of the CICR across the entire SR, the experimentally observed threshold (figure 1) would be much lower and the rate of junctional coupling by CICR faster. In other words, due to the high [Ca^2+^]_i_ threshold of about 10 *µ*M for Ca^2+^ activation of RyR3, CICR engendered by a single L-SR junction is likely to die out, unless reinforced by a process of quantal releases by other L-SR junctions within a cluster [65]. Therefore it seems more likely that multiple L-SR junctions work in concert in a process characterized by both additive and regenerative elements to provide the necessary threshold and margin of safety required to ensure the propagation by CICR of the global wave via the more widely distributed RyR2 along extra-junctional SR, once the Ca^2+^ release wave escapes an RyR3-enriched SR region, as previously suggested in [11].

In this respect, our results also allowed us to establish a time interval for the duration of the transient (*t*_transient_) that would be compatible with the hypothesis that sequential summation of all-or-none lysosomal Ca^2+^ release events from a set of individual L-SR junctions is responsible for generating Δ[Ca^2+^]_shoulder_. Remarkably, this value is comparable to the average duration of the simulated transients, which was determined on the basis of ultra-structural details and completely independently of the summation effect hypothesis. The assumption of summation may imply that the role of the NAADP as a stimulus is limited to the initial lysosomal Ca^2+^ release, while the recruitment of subsequent L-SR junctions may be governed by further lysosomal calcium release, by SR Ca^2+^ release via Ca^2+^ activated junctional RyR3, and/or propagation and combination of calcium signals via inter-junctional clusters of RyR3. Therefore, from our model we envision the intriguing possibility of an important regulatory role of lysosomal Ca^2+^ content by SR Ca^2+^, in such a way that summation of calcium signals at multiple junctional complexes may give rise to the shoulder. While only further experiments can verify this, the quantitative corroboration provided by our findings sets on firmer ground the conclusion from earlier studies that Ca^2+^ bursts from lysosomes are indeed responsible for initiating the first phase of a cell wide Ca^2+^ wave via L-SR junctions.

### 3.4 Disruption of nanojunctions, plasticity and pathology

Carrying the signal time-scale analysis further and using the calculated *t*_transient_ to take a temporal average of the simulated [Ca^2+^]_NJ_ at different junctional widths, we find that above a width of about 30 nm the transients would be unable to trigger Ca^2+^ release from the RyR3s (figure 7B). Thus it is possible that heterogeneity and plasticity are controlled by a variable width of the junctional nanospace. For example, in atrial myocytes it has been proposed that NAADP evokes Ca^2+^ release from an acidic store, which enhances general SR Ca^2+^ release by increasing SR Ca^2+^ load and activating RyR sites [34]. This functional variant may be provided by either: (1) An increase in junctional distance such that [Ca^2+^]_NJ_ is insufficient to breach the threshold for activation of RyR2, yet sufficient to allow for increases in luminal Ca^2+^ load of the SR via apposing SERCA2 clusters; or (2) L-SR junctions in cardiac muscle formed between lysosome membranes and closely apposed regions of the SR which possess dense SERCA2 clusters and are devoid of RyR2. Further ultrastructural studies on cardiac muscle and other cell types may therefore provide a greater understanding of how L-SR junctions may have evolved to provide for cell-specific modalities within the calcium signaling machinery.

Under metabolic stresses, such as hypoxia, lysosomal pathways have been proposed to provide for autophagic glycogen metabolism via acid maltase in support of energy supply, and significant levels of protein breakdown during more prolonged metabolic stress. Moreover, although controversial, in some cell types it has also been suggested that lysosomes may contribute to the energy supply by providing free fatty acids for beta-oxidation by mitochondria [66]. It may be significant, therefore, that TPC2 gating and thus autophagy may be modulated by mTOR [31, 67]. This may well speak to further roles for lysosomal calcium signaling and L-SR junctional plasticity during hypoxic pulmonary vasoconstriction and even during the development of hypoxia pulmonary hypertension [68], not least because AMP-activated protein kinase has been shown to modulate autophagy through the phosphorylation and inhibition of mTOR (see for example, [69]).

The process of autophagy serves not only to regulate programmed cell death, but also to recycle organelles, such as mitochondria, through a process of degradation involving lysosomal hydrolases [70]. For this to occur L-SR junctions would likely be disrupted and thus select for local rather than global Ca^2+^ signals in order to facilitate fusion events among lysosomes, endosomes, autophagosomes and amphisomes [71], which follows from the fact that Ca^2+^ plays a pivotal role in vesicle trafficking and fusion [72]. Active non-synchronous movements of TPC2-expressing vesicles have been detected in live-cell imaging experiments using GFP-tagged proteins [20]. Thus, lysosomal Ca^2+^ signaling may modulate plasticity in the manner required at all stages of multiple membrane fusion events that are dependent on Ca^2+^ for the effective formation of the SNARE complexes [72]. In short, transport of proteins between lysosomes, Golgi apparatus, and plasma membrane via lysosomes and endosomes may be coordinated at different stages by both global Ca^2+^ signals and spatially restricted Ca^2+^ release from acidic stores in a manner that is in some way determined by L-SR junctional integrity.

Loss of integrity of L-SR junctions may also contribute to disease, given that lysosomal Ca^2+^ release both depletes luminal Ca^2+^ and causes intraluminal alkalinization [73]. Therefore, disruption of L-SR junctions may modulate the activity of pH-sensitive hydrolytic lysosomal enzymes, such as glucocerebrosidase and acid sphingomyelinase, which exhibit a marked loss of function at pH *>* 5 [21, 74] that could lead to accumulation of macro-molecules such as glucocerebroside and sphigomylin. As mentioned previously, dysfunction of these enzyme systems consequent to L-SR junctional abnormalities could also contribute to pathologies associated with subclasses of lysosome storage disease such as Niemann-Pick disease type C1 [22, 23], Pompe and Gaucher [24, 25], which may include hepatic portal [23], or pulmonary hypertension [24,25], dysfunctions in cholesterol trafficking [26] and consequent increases in plasma cholesterol levels, vascular lesion formation, atherosclerosis/thrombosis and medial degradation [23, 27].

### 3.5 Conclusion

We have determined that L-SR junctions, about 15 nm in width and extending to ≈ 300 nm in lateral dimensions are a regularly occurring feature in rat pulmonary arterial smooth muscle cells, in which L-SR junctions were first proposed. The present study provides a mechanistic basis for the observed NAADP-induced Ca^2+^ signals and strong support for the proposal that these Ca^2+^ signals are generated at L-SR junctions. Even within the variability of our recorded values of L-SR junctional widths and extension, our results suggest that localized [Ca^2+^] transients due to junctional Ca^2+^ release can without fail reach values required to breach the threshold for CICR from junctional RyR3s. Perhaps most significantly, disruption of the nanojunctions decreases [Ca^2+^]_NJ_ below the value for CICR via junctional RyR3s. Therefore, consistent with previous studies on the PM-SR membrane [39], we have established that L-SR junctions are required to allow such signals to be generated and that there is a 30–50 nm limit on junctional width, above which there is loss of junctional integrity and inadequate control of ion movements within the junctional space. This suggests that the observed L-SR junctions in PAMSCs are not only capable of delivering localized Ca^2+^ bursts of the required magnitude, but are also necessary if lysosomes are to fulfill this identified role in Ca^2+^ signaling. In other words, L-SR nanojunctions are a necessary and sufficient condition for generating local Ca^2+^ bursts essential for NAADP-induced Ca^2+^ waves. Most importantly, however, this study demonstrates the importance of junctional architecture on the nanoscale to the capacity for coupling across cytoplasmic nanospaces, tight regulation of ion transport and thus signal transduction. In turn, this highlights the possibility that alterations in the dimensions and architecture of intracellular nanojunctions lead to cell dysfunction and hence disease.

## 4 Materials and methods

All the experiments and procedures were carried out in accordance with the guidelines of the University of British Columbia Animal Care Committee and in accordance with the United Kingdom Animals (Scientific Procedures) Act 1986.

### 4.1 Electron microscopy

Male Wistar rats weighing 220–250 g were anesthetized with 3 mL of Thiotal. The thoracic cavity was opened and flooded with warm physiological saline solution (PSS) containing 145 mM NaCl, 4 mM KCl, 1 mM MgCl_2_, 10 mM HEPES, 0.05 mM CaCl_2_, and 10 mM glucose (pH 7.4). After extraction of the heart and lungs and their placement in HEPES buffer, rings from the primary and secondary branches of the pulmonary artery were dissected and immediately immersed in fixative solution. The primary fixative solution contained 2.5% glutaraldehyde in 0.1 M sodium cacodylate buffer at room temperature. The artery rings were then washed three times in 0.1 M sodium cacodylate (30 min in total). In the process of secondary fixation, the tissue rings were fixed with 1% OsO_4_ in 0.1 M sodium cacodylate buffer for 1 h followed by three 10-minute washes with distilled water and *en bloc* staining with 2% uranyl acetate. The samples were then dehydrated in increasing concentrations of ethanol (25, 50, 75, 80, 90, and 95%). In the final process of dehydration, the samples underwent 3 washes in 100% ethanol. The artery rings were then resin-infiltrated in increasing concentrations (30, 50, and 75% in ethanol) of a 1:1 mix of Epon and Spurr’s resins. The infiltration process was completed by three passages in 100% resin. All of the ethanol dehydration and resin infiltration steps were carried out by using a laboratory microwave oven. The blocks were finally resin-embedded in molds and polymerized overnight in an oven at 60°C.

#### 4.1.1 2D imaging

For standard (2D) electron microscopy imaging, 80-nm sections were cut from the embedded sample blocks on a Reichert Ultracut-E microtome using a diamond knife and were collected on uncoated 100- and 200-mesh copper grids (hexagonal or square meshes). The sections were post-stained with 1% uranyl acetate and Reynolds lead citrate for 12 and 6 minutes, respectively. Electron micrographs at various magnifications were obtained with a Hitachi 7600 transmission electron microscope at 80 kV.

Lysosomes in these images were identified according to their well established appearance features: they are single lipid bilayer membrane-bound, with a granular, more or less uniform luminal matrix that is more electron dense than the surrounding cytosol. Secondary lysosomes may also contain less granular structures within the finer matrix. Moreover, lysosomes are normally distinguishable from endosomes by their larger size, hence we set a threshold “diameter” of *>* 200 nm for acceptance of a lysosome, below which all vesicles were excluded.

#### 4.1.2 Tomography (3D imaging)

To obtain electron microscopic tomograms, we cut 200-nm-thick sections from the same sample blocks used for standard imaging. The sections were then collected on Formvar coated slot copper grids and post-stained with 1% uranyl acetate and Reynolds lead citrate for 20 and 10 minutes, respectively. We surveyed the sample sections using a FEI Tecnai G2 200 kV transmission electron microscope and identified regions of interest containing L-SR junctions. We then acquired tomograms of several of those regions by taking 2D scans through the sample every 5° of inclination as it was tilted between −60° and +60° with respect to horizontal. The scans were reconstructed with the Inspect3D software tools and structures of interest, primarily lysosomal and SR membranes in the same cellular neighbourhood, were traced out using Amira software.

### 4.2 Image and data analysis

The images of the samples were further processed using GIMP (GNU Imaging Manipulation Program, open source, available at gimp.org) to enhance membrane contrast in the interest of improving our characterization of the L-SR junctions.

The SR and lysosomal membranes were outlined, highlighted and measured in pixels using the Inkscape software (open source, available at inkscape.org), converting the pixel measurements to nm using the scale bar in the recorded micrographs. By modifying the Inkscape script for measuring lengths, we were able to output the measurements directly into a text file, which we used to produce the histograms in Figure 3B–D. We used the package Gnuplot (open source, available at gnuplot.info) to produce the histograms and plots presented herein.

### 4.3 Ca^2+^ imaging

Single arterial smooth muscle cells were isolated from second-order branches of the pulmonary artery. Briefly, arteries were dissected out and placed in low Ca^2+^ solution of the following composition (mM): 124 NaCl, 5 KCl, 1 MgCl_2_, 0.5 NaH_2_PO_4_, 0.5 KH_2_PO, 15 NaHCO_3_, 0.16 CaCl_2_, 0.5 EDTA, 10 glucose, 10 taurine and 10 Hepes, pH 7.4. After 10 min the arteries were placed in the same solution containing 0.5 mg/ml papain and 1 mg/ml bovine serum albumin and kept at 4°C overnight. The following day 0.2 mM 1,4-dithio-DL-threitol was added to the solution, to activate the protease, and the preparation was incubated for 1 h at room temperature (22°C). The tissue was then washed at 3°C in fresh low Ca^2+^ solution without enzymes, and single smooth muscle cells were isolated by gentle trituration with a fire-polished Pasteur pipette. Cells were stored in suspension at 4°C until required.

PASMCs were incubated for 30 min with 5 *µ*M Fura-2-AM in Ca^2+^-free PSS in an experimental chamber on a Leica DMIRBE inverted microscope and then superfused with Fura-2 free PSS for at least 30 min prior to experimentation. Intracellular Ca^2+^ concentration was reported by Fura-2 fluorescence ratio (F340/F380 excitation; emission 510 nm). Emitted fluorescence was recorded at 22°C with a sampling frequency of 0.5 Hz, using a Hamamatsu 4880 CCD camera via a Zeiss Fluar 40×, 1.3 n.a. oil immersion lens and Leica DMIRBE microscope. Background subtraction was performed on-line. Analysis was done via Openlab imaging software (Improvision, UK).

NAADP was applied intracellularly in the whole-cell configuration of the patch-clamp technique, and in current clamp mode (*I* = 0) as described previously [17]. The pipette solution contained (in mM): 140 KCl, 10 Hepes, 1 MgCl_2_ and 5 *µ*M Fura-2, pH 7.4. The seal resistance, as measured using an Axopatch 200B amplifier (Axon Instruments, Foster City, CA), was ≥ 3 GΩ throughout each experiment. Series resistance and pipette resistance were ≤ 10 MΩ and ≤ 3 MΩ, respectively. All experiments were carried out at room temperature (*≈* 22° C).

### 4.4 Quantitative modeling

The main stages of the quantitative modeling approach are:

1. the design of 3D software mesh objects (nets of interconnected triangles by which surfaces can be represented in computer graphics) representing a typical L-SR region, including a whole lysosome and a portion of neighbouring SR network. These objects are built to-scale following the ultrastructural characterization of the L-SR junctional regions as it results from our electron microscopy image analysis; this phase was carried out using the “3D content creation suite” Blender (open source, available at blender.org);
2. the positioning of the relevant transporters on the reconstructed membranes (TPC2 complexes on the lysosome, SERCA2a and RyR3 on the SR) according to information gathered from the literature on their typical membrane densities and the implementation of the transporter known kinetics and multistate models and of the ion diffusivities (Ca^2+^ and mobile Ca^2+^ buffers);
3. the simulation of molecular Brownian motion in the cytosol by random walk algorithms; this phase was performed by writing appropriate code for the stochastic particle simulator MCell (freely available at mcell.org) [35–37]. In a nutshell, MCell reproduces the randomness of the molecular trajectories, of the ion transporter flickering and of the relevant chemical reactions by probabilistic algorithms governed by random number generators (iterative mathematical algorithms, which produce a random sequence of numbers once initiated by a given number called seed). This enables the simulation of a number of microphysiological processes, all stochastically different from one another. The average outcome of the processes, to mimic the instrumental output during experimental measurements, is obtained by taking the average of a desired quantity, e.g., [Ca^2+^], over a large number of simulations all initiated with a different seed;
4. the measurement of simulated [Ca^2+^] in the L-SR junctions from the process of Ca^2+^ release via TPC2-related signaling complex on the lysosome and SR Ca^2+^ uptake by the SERCA2a pumps, and static as well as dynamic visualization of the simulations; this stage is part of MCell’s output.

## Acknowledgements

We are very grateful to Garnet Martens and the University of British Columbia Bioimaging Facilty for their assistance. We also acknowledge the help of David Walker and Arash Tehrani with TEM image analysis and acquisition. This research has been enabled by the use of computing resources provided by WestGrid and Compute/Calcul Canada (westgrid.ca, computecanada.ca).

